# Five hours total sleep deprivation does not affect CA1 dendritic length or spine density

**DOI:** 10.1101/2021.02.04.429550

**Authors:** Alvin T.S. Brodin, Sarolta Gabulya, Katrin Wellfelt, Tobias E. Karlsson

## Abstract

Sleep is essential for long term memory function. However the neuroanatomical consequences of sleep loss are disputed. Sleep deprivation has been reported to cause both decreases and increases of dendritic spine density. Here we use Thy1-GFP expressing transgenic mice to investigate the effects of acute sleep deprivation on the dendritic architecture of hippocampal CA1 pyramidal neurons. We found that five hours of sleep deprivation had no effect on either dendritic length or dendritic spine density. Our work suggests that no major neuroanatomical changes result from one episode of sleep deprivation.

## Introduction

Sleep disturbance is a feature of several prevalent brain diseases, including Alzheimer’s disease, anxiety, depression and schizophrenia (Wulff et al., 2010), and even our distant ancestors were plagued by poor sleep (Ancoli-Israel, 2001). There is widespread concern that our modern lifestyle is making sleep disorders more prevalent. Lifestyle factors such as altered working patterns and use of electronic devices have been proposed to fuel an epidemic of sleep deprivation (Shochat, 2012). However whether modern humans truly sleep less is debated (Yetish et al., 2015).

Adequate sleep is essential to cognitive function across the animal kingdom (Keene and Duboue, 2018), and typically consists of both rapid eye movement sleep (REM-sleep) and slow-wave sleep (SWS) (Cousins and Fernández, 2019). The homeostatic necessity of SWS seems to be stronger than that of REM-sleep, as the proportion of SWS increases after sleep deprivation (Dijk, 2009; Rodriguez et al., 2016). In healthy humans, the first half of a night’s sleep is dominated by SWS with increasing proportions of REM-sleep observed during the latter half.

Particularly strong, and particularly studied, is the association between sleep and memory function (Cousins and Fernández, 2019). Sleep prior to learning is necessary for proper encoding. Extended periods of wakefulness impair attention (Krause et al., 2017; Lim and Dinges, 2010), and induction of long-term potentiation is impaired after sleep deprivation (Campbell et al., 2002). Encoding seems to be reliant on SWS, as selectively disturbing SWS disrupts encoding the following day (Van Der Werf et al., 2009). The role of REM-sleep preceding learning is less clear, with one study reporting that disturbing REM has no effect on subsequent memory encoding (Kaida et al., 2015). Sleep following learning is also essential, as learning events followed by sleep are retained better than those followed by a period of wake (Igloi et al., 2015).

The mechanism behind the necessity and behavioural effects of sleep are not fully understood. One influential theory for the functional role of sleep is the synaptic homeostasis hypothesis (SHY). SHY states that experiences during the day are encoded by increases in synaptic weight, necessitating homeostatic downscaling during sleep to prevent overexcitability (Tononi and Cirelli, 2014). Reduced memory function would result from impaired consolidation when sleep is disturbed, as well as from prolonged wakefulness causing general overexcitability that impairs signal recognition and hinders further potentiation (Tononi and Cirelli, 2014).

A key prediction by SHY is that global synaptic strength should increase during periods of wake and decrease during sleep, as opposed to homeostatic downscaling occurring continuously. This prediction has been tested in several ways. Sleep deprivation has been found to lower seizure thresholds in both humans (Malow, 2004) and animal models (Giorgi et al., 2014). However, decreased firing thresholds have only been demonstrated in epileptic patients, not healthy subjects (Giorgi et al., 2014). At the structural level, *in vivo* two-photon imaging has demonstrated an average decrease of spine size and GluA1 content during sleep, with a small subset of spines instead growing and increasing their GluA1 content (Diering et al., 2017). Phosphorylation of several plasticity-related proteins also vary in a manner suggesting synaptic potentiation during wake and downscaling during sleep (Vyazovskiy et al., 2008).

Histological studies of hippocampus offer conflicting evidence with regards to how sleep deprivation affects synapse numbers and dendritic structure. Some studies show that sleep deprivation increases spine density in CA1 as predicted by SHY (Gisabella et al., 2020). However, other studies show that sleep deprivation causes a loss of spines in CA1 (Havekes et al., 2016) and the dentate gyrus (Raven et al., 2019), even shortening the dendritic tree as a whole in CA1 (Havekes et al., 2016). Should sleep deprivation cause the dendritic arbour to shrink and reduce synapse numbers, rather than the opposite, this would be a strong argument against SHY and necessitate looking for alternate explanations regarding the function of sleep (Raven et al., 2018). In this study we utilise recent innovations of tissue imaging to determine the effects of sleep deprivation on dendritic arbour and spine density in the CA1 of hippocampus.

## Methods

### Animals

Male Thy1-GFP line M mice obtained from Jackson aged 12 to 44 weeks were used for all experiments. Mice were housed in a 12h light cycle with lights on at 7 AM and food and water ad libitum. Animals were split into the three groups based on cages in a pseudorandom schedule.

### Sleep Deprivation

Animals were kept in their home cages with 4-6 animals per cage for the duration of the experiments. Mice were handled daily for 3 days prior to the experiment. They were sleep deprived from 7 AM to 12 AM using the gentle handling method (Colavito et al., 2013), conducted by two experimenters. When mice were about to fall asleep they were awoken by, in ascending order; gently tapping the cage, gently tilting the cage, or disturbing the bedding. Control mice, and mice undergoing recovery sleep, were kept in the same room as the sleep deprived mice.

### Tissue Processing

At the end of the experiment mice were anesthetised with pentobarbital and perfused with 4% formalin in phosphate buffered saline (PBS). Whole brains were removed and post-fixed in 4% formalin for 24 hours, and subsequently stored in PBS.

Brains were embedded in an agarose gel and sectioned by vibratome into 300 µm coronal sections for dendrite tracing and 200 μm coronal sections for spine analysis.

The sections were cleared using an adapted SeeDB2 protocol (Ke et al., 2016). Briefly, brain tissue was incubated in serial solutions of 2% Saponin with increasing concentrations of Omnipaque 350.

### Imaging

For dendrite tracing, images were acquired using a Zeiss Lightsheet Z.1 microscope (Light Sheet Microscopy Pilot Facility, KTH), equipped with a 20x/1.0 NA objective and excitation was performed by two LSFM 10x/0.2NA illumination objectives.

For spine analysis, tissue was stained using an anti-GFP alexa 555 antibody to enhance signal.

Images were acquired with an Axioobserver Z1 microscope (Carl Zeiss, Oberkochen, Germany, provided by Biomediucm Imaging Core facility), with a 63x oil (NA 1.4) with a metal halide mercury light as a light source, with an excitation wavelength of 553 nm and emission 568 nm. Z stacks of apical dendrites were obtained with a pixel resolution of 0.04 μm in XY and a 0.2 μm interval between Z slices.

### Dendritic Tree Analysis

The lightsheet image stacks were stitched together using manual alignment in Arivis Vision4D. Apical dendrites of the CA1 pyramidal neurons were traced by a blinded investigator using the Autopath semiautomated tracing of Imaris version 9.6.0(Filament tracker licence, Imaris).

### Statistical Analysis

Statistical analysis was performed using R, version 3.6.3, with RStudio, version 1.2.5033.

A mixed effects linear model was constructed using the R package lme4 version 1.1-23, with the specified dendritic parameter given as a function of treatment, with mouse identity as a random effects factor. Significance testing of treatment effects was performed using the R package lmerTest v. 3.1-3. A p-value of less than 0.05 was considered significant.

Post-hoc power analysis was performed by Monte Carlo simulations using the R package Simr version 1.0.5. For the simulations, expected effect size of 15% or 30% were used, corresponding to the range of effect sizes seen in the literature.

## Results

### Sleep deprivation does not alter the shape of the dendritic tree

We first set out to investigate the effects of sleep deprivation on dendritic length. Thy1-GFP mice were subjected to one of the following treatments: sleep deprivation for 5 h (n=10 mice), no sleep deprivation (n=12 mice), or 5 h of sleep deprivation followed by 3 h of recovery sleep (n=8 mice) (Fig 1A). Using light-sheet imaging and semi-automated tracing, apical dendrites from 7-8 CA1 pyramidal neurons were traced from each animal (Fig 1B).

**Figure 1.**
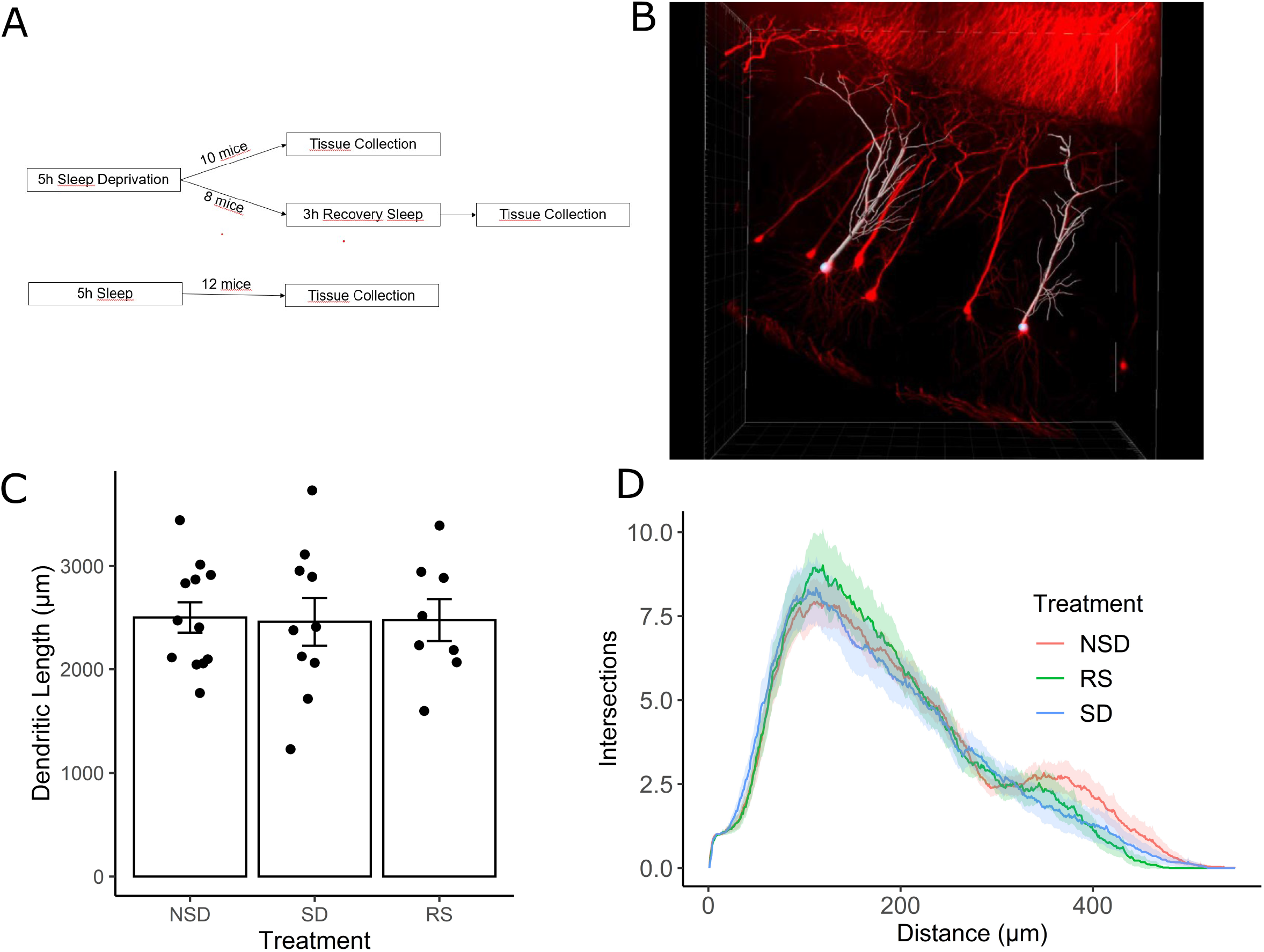
Sleep deprivation does not impact dendritic length ofhippocampal CAI neurons. (A) Experimental outline. (B) Representative image ofThyl-GFP CAI neurons (red) and semi-automated tracing using IMARIS (white). (C) Shows the average length ofapical dendrites per treatment: 5h sleep deprivation (SD, n=lO mice) or SD followed by 3h recovery sleep (RS, n=8 mice) or no sleep deprivation (NSD, n= 12 mice). (D) Averaged sholl diagram ofdendrites per treatment. The shaded area denotes the 95% confidence interval.

We found no significant effect on overall dendritic length by sleep deprivation (p=0.847), or recovery sleep (p=0.923). It is possible that sleep deprivation does not significantly alter the overall length of the dendritic tree, but instead affects a particular part of the dendritic tree. To investigate this possibility we performed Sholl analysis of the traced neurons, and found that the shapes of the Sholl diagrams did not differ significantly between treatments (Fig 1D).

Monte Carlo simulations were used to perform a post-hoc power analysis based on our model. We found that our study had a 30% (95% CI 26.88% – 32.64%) chance to detect a 15% (Cohen’s d 0.5) difference in dendritic length comparing sleep deprivation vs no sleep deprivation, or a 76% (95% CI 73.02-78.43) chance to detect a 30% difference (Cohen’s d 1.0), corresponding to the effect size detected reported by Havekes et al (Havekes et al., 2016).

### Sleep deprivation does not have major effects on dendritic spine density

We next analyzed changes in the dendritic spines of CA1 neurons. Dendritic spines have been shown to be more motile structures than dendritic branches, and thus possibly more susceptible to sleep deprivation. Using tissue from the same mice as for the dendritic length analysis, tissue was imaged using an LSM-800 airyscan confocal microscope. Dendritic spines were counted on 3rd to 5th order apical dendrites of CA1 pyramidal neurons from sleep deprived (n=6 mice), recovery sleep (n=4 mice) or non sleep deprived (n=6) mice (Fig 2A). Spines were counted on an average of 6.3 dendrites per mouse.

**Figure 2.**
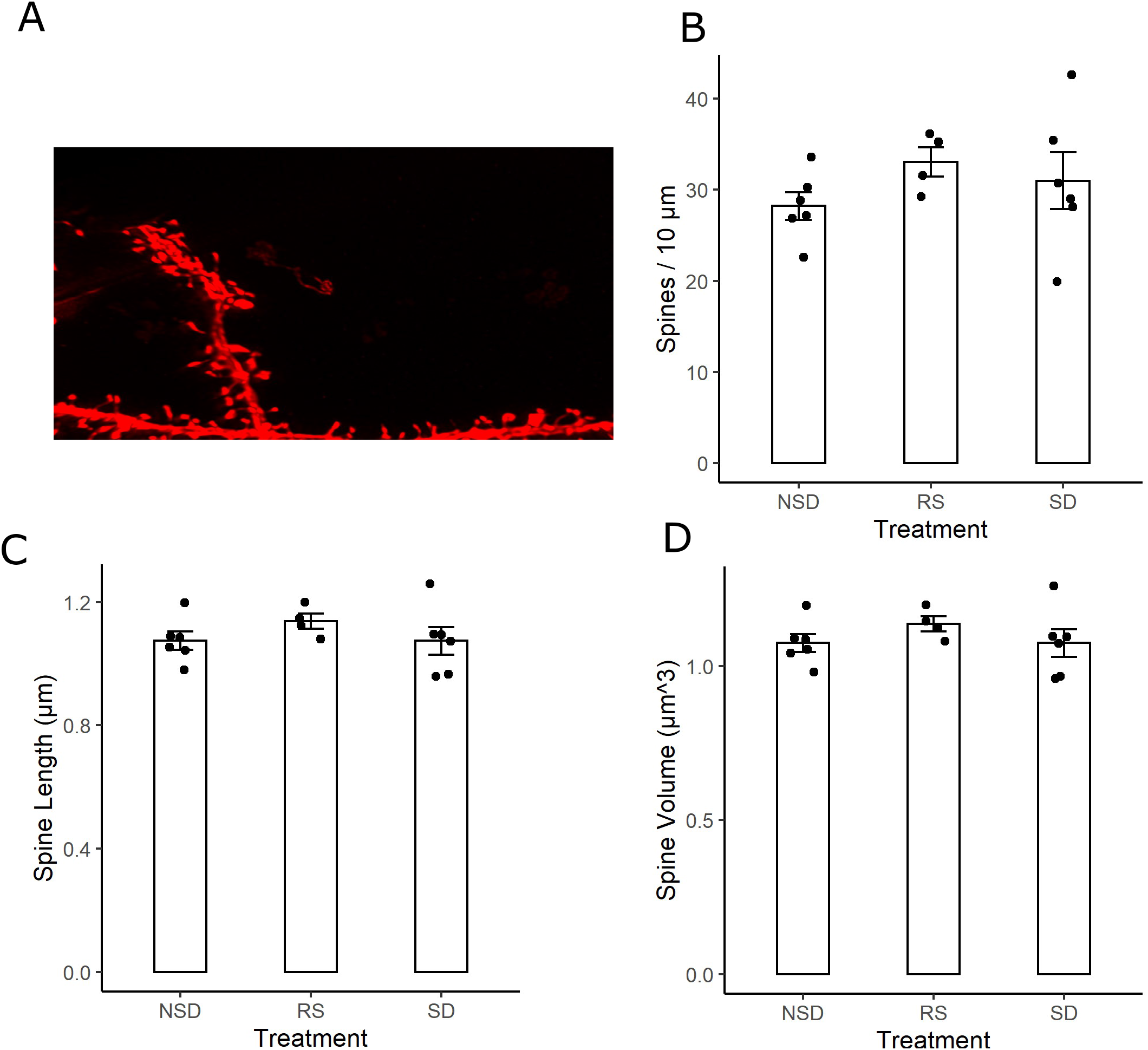
No detectable differences in spine density or morphology ofhippocampal CAI neurons after sleep deprivation. (A) Representative picture ofThyl-GFP third-order dendritic branch used for spine counting. Scale bar 5 μm. (B) Shows the average total spine density for; 5h sleep deprivation (SD, n=6 mice), SD followed by 3h recovery sleep (RS, n=4 mice) or no sleep deprivation (NSD, n=6 mice). (C) Shows the average spine length ofthe measured spines. (D) Shows the average volume of the spines.

There were no significant effects of sleep deprivation on dendritic spine density (Fig 2B). Compared to non-sleep deprived mice there was a non-significant higher mean spine density in sleep deprived mice (p=0.464) and recovery sleep mice (p=0.211) (Fig 2B). The shape of dendritic spines were unchanged, with no significant effects by any treatment on the average length of dendritic spines (Fig 2C) or the volume of dendritic spines (Fig 2D). There was no difference in dendrite diameter across the groups (Fig S1A). The branch order of analysed neurons was similar across groups (Fig S1B), and spine density did not vary significantly as a function of branch order (Fig S1C).

We also performed a post-hoc power analysis on this experiment. The tissue from several mice were lost during processing for spine analyisis. Hence, the number of animals in this part of the study is lower than for the dendritic tracing. We estimated that we had 25% (95% CI 22.25-27.70) power to detect a 15% (Glass’s delta 0.85) difference in dendritic spine density between sleep deprived and not sleep deprived mice, or a 63% (95% CI 60.13-66.20) power to detect a 30% difference (Glass’s delta 1.69). This can be put into contrast of the previously reported differences that has been in the range of 15-30% (Gisabella et al., 2020; Havekes et al., 2016).

## Discussion

That adequate sleep is vital for the formation of lasting memories has been clearly established (Cousins and Fernández, 2019), but the mechanisms involved remain disputed. Charting the neuroanatomical changes caused by sleep deprivation promises to offer clues to its function, but conflicting evidence points to both synapse loss (Havekes et al., 2016; Raven et al., 2019) and synapse gain during sleep deprivation (Gisabella et al., 2020). Dramatically, it has even been reported that brief sleep deprivation can shorten the dendritic tree as a whole in CA1 pyramidal cells (Havekes et al., 2016).

Here, we show that no major changes occur in the dendritic tree of CA1 neurons after brief (5 hours) sleep deprivation. There were no significant changes of overall dendritic length nor in the shape of the dendritic arbour as shown by Sholl analysis. There are several methodological differences that could explain the discrepancy between our results and those of Havekes et al, where a substantial reduction of dendritic length after sleep deprivation was found. Where they used Golgi staining, we instead used transgenic Thy1-GFP mice. The average apical dendrite of our control group was traced to 2500 μm, while the control neurons’ apical dendrites in Havekes et al were 1200 μm. This could indicate that our GFP-labelled neurons were more completely traced, which could be one reason for the discrepant results.

Differences in statistical methods could also lead to discrepancies in results. A common practise in neuroanatomical studies is to treat studied neurons as one homogenous group with regards to statistical analysis. However, disregarding the inter-relatedness of neurons from the same animal risks underestimating the variance in the sample and leading to too small sample sizes being used (Wilson et al., 2017). A mixed effects model avoids this problem, but the statistical tests used specifically for the neuroanatomical analysis have unfortunately not been listed in previous studies on the topic. To our knowledge, ours is the best powered study to date on the effect of sleep deprivation on dendritic length. The small size of both studies leaves open the possibility that different results could have been obtained by chance, without the true effect differing between the studies.

Studies using *in vivo* transcranial imaging support the view of a largely static dendritic tree. Individual dendritic branches have been imaged successfully over several weeks (Holtmaat and Svoboda, 2009), a feat which would be impossible if large-scale remodelling of the dendritic arbour was commonplace. However, these experiments did not feature sleep deprivation, and cannot rule out dendritic changes occurring specifically in this setting.

Our findings on dendritic spines were less conclusive than those on dendritic length, due to a loss of tissue from several mice prior to this step impairing statistical power. No changes in spine number or morphology were found, a non-significant increases of spine density after sleep deprivation was noted. Only tentative conclusions can be drawn, with the data weakly speaking against large decreases in spine number after sleep deprivation.

The dynamics of dendritic spines during sleep have been more extensively studied than those of the dendritic tree as a whole. Several studies have found that spines are pruned during sleep (de Vivo et al., 2017; Diering et al., 2017; Li et al., 2017, 2017; Vyazovskiy et al., 2008). This has been found to occur during both SWS (Feld and Born, 2017) and REM sleep (Li et al., 2017). Newly formed spines are selectively maintained during this process, resulting in a net loss of spines while consolidation of strong synapses occurs (Li et al., 2017). *In vivo* imaging thus provides support for the idea that the net effect of sleep is a pruning of synaptic spines, which agrees with some histological studies (Gisabella et al., 2020; Spano et al., 2019) but not others (Havekes et al., 2016; Raven et al., 2019).

Notably, sleep deprivation entails not only the absence of sleep, but also an abnormally long period of wake, which may have its own effects. In addition, all sleep deprivation methods are associated with some level of stress (Nollet et al., 2020). Gentle handling was chosen as the method of sleep deprivation in the present study as it is less stressful than many other methods (Nollet et al., 2020), but not stress-free. The method is hard to standardize between experimenters, and inter-experimenter differences in odour, training and demeanour could all affect the stress levels of the sleep deprived animals. Indeed, stress has been shown to adversely affect dendritic spine density (Leuner and Shors, 2013), which might explain why studies may differ with respect to dendritic spine density.

Our study shifts the balance of evidence away from reductions of the dendritic tree occurring during sleep deprivation, and to a lesser extent away from reductions of spine density. This weighs in favour of the synaptic homeostasis hypothesis, which is largely incompatible with sleep deprivation causing a decrease in synapse number. This is an important issue with implications for our understanding of the role of sleep as a whole, but before final conclusions can be drawn there is an urgent need for further, statistically robust studies on this topic.

## Acknowledgements

We would like to thank the Rut och Arvid Wolffs minnesstiftelse, Vetenskapsrådet, Hjärnfonden, Per Nydahl and the Karolinska institute with the funding programs KID and CSTP for supporting this work. We would like to thank the Lightsheet pilot facility for providing the lightsheet microscope and for excellent technical support.

---

**Figure S1.**
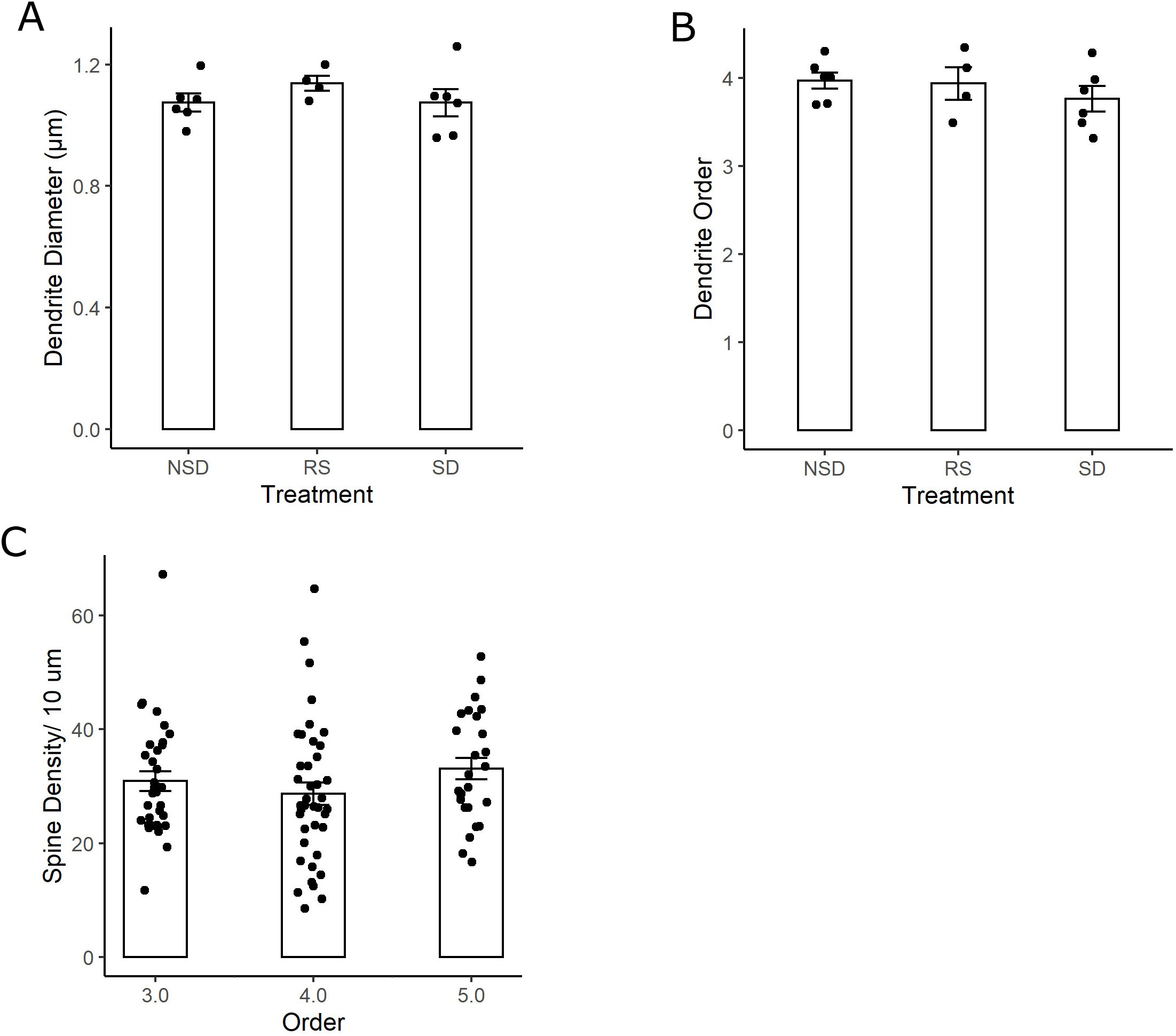
Comparable dendritic branches analysed across treatments. (A) No differences in dendrite diameter between mice subjected to 5h sleep deprivation (SD, n=6 mice), sleep deprivation followed by 3h recovery sleep (RS, n=4 mice), or no sleep deprivation (NSD, n=6 mice). (B) Similar ordered dendrites imaged across all three treatments. (C) No difference in spine density across analysed dendrite branch orders.

